# The pathognomonic FOXL2 C134W mutation alters DNA binding specificity

**DOI:** 10.1101/2020.03.20.984476

**Authors:** Annaïck Carles, Genny Trigo-Gonzalez, Rachelle Cao, S.-W. Grace Cheng, Michelle Moksa, Misha Bilenky, David G. Huntsman, Gregg B. Morin, Martin Hirst

**Affiliations:** Department of Microbiology and Immunology, Michael Smith Laboratories, University of British Columbia, Vancouver, British Columbia, V6T1Z4, Canada; Canada’s Michael Smith Genome Sciences Centre, BC Cancer, Vancouver, British Columbia, V5Z1L3, Canada; Department of Molecular Oncology, BC Cancer, Vancouver, British Columbia, V5Z 4E6, Canada; Department of Medical Genetics, University of British Columbia, Vancouver, British Columbia, V6H 3N1, Canada; Department of Pathology and Laboratory Medicine, University of British Columbia, Vancouver, British Columbia, V6T 2B5, Canada; Department of Obstetrics and Gynaecology, University of British Columbia, Vancouver, British Columbia, V6Z 2K8, Canada

**Author notes:** Correspondence (M.H.) and (G.B.M.).

## Abstract

The somatic missense point mutation c.402C>G (p.C134W) in the FOXL2 transcription factor is pathognomonic for adult-type granulosa cell tumours (AGCT) and a diagnostic marker for this tumour type. However, the molecular consequences of this mutation and its contribution to the mechanisms of AGCT pathogenesis remain unclear. To explore the mechanisms driving FOXL2^C134W^ pathogenicity we engineered V5-FOXL2^WT^ and V5-FOXL2^C134W^ inducible isogenic cell lines and performed ChIP-seq and transcriptome profiling. We found that FOXL2^C134W^ associates with the majority of the FOXL2 ^WT^ DNA elements as well as a large collection of unique elements genome-wide. We confirmed an altered DNA binding specificity for FOXL2^C134W^ *in vitro* and identified unique targets of FOXL2^C134W^ including *SLC35F2* whose expression increased sensitivity to YM155 in our model.

**Statement of Significance:** Mechanistic understanding of FOXL2^C134W^ induced regulatory state alterations drives discovery of a rationally designed therapeutic strategy.

## Introduction

Adult Granulosa Cell Tumours (AGCTs) of the ovary are sex cord-gonadal stromal tumours that are thought to be derived from granulosa cells (GCs). GCs form a multilayered cluster surrounding the oocyte of ovarian follicles and proliferate throughout folliculogenesis to support oocyte maturation. AGCTs are uncommon tumours accounting for about 5% of all ovarian cancers^1^. The somatic missense point mutation c.402C>G (p.C134W) in FOXL2 is pathognomonic for AGCTs (Shah et al. 2009) providing a specific diagnostic marker for this disease^2-4^.

FOXL2 is essential for normal ovary development and is a member of the large Forkhead Box (FOX) family of transcription factors that are characterized by the presence of a highly conserved winged helix/forkhead DNA-binding domain^5^. The mutation at amino acid position 134 of FOXL2, replacing a highly conserved cysteine with tryptophan, is located in the second wing structure at the edge of the DNA binding domain and is adjacent to the SMAD3 dimerization domain^6,7^. Unlike most FOX transcription factors, which bind DNA as monomers^8^, FOXL2 associates with DNA as either a homodimer^9^ or heterodimer with partners such as SMAD3^10,11^. Structural studies suggest the C134W mutation does not affect FOXL2 protein folding or DNA binding but rather its interaction with partners^7^. Functional studies suggest it leads to perturbation of steroidogenesis, apoptosis, and cell proliferation signaling pathways^10-14^ including TGF-β signaling^15^, the target of a recent phase I clinical study^16^. However, direct targets of FOXL2^C134W^ and the mechanisms by which this mutation alters FOXL2-mediated transcriptional regulation remain elusive.

To define direct transcriptional targets of FOXL2^C134W^, we engineered V5-FOXL2^WT^ and V5-FOXL2^C134W^ inducible isogenic cell lines and performed chromatin immunoprecipitation sequencing (ChIP-seq) and RNA sequencing (RNA-seq) following induction of the V5 tagged FOXL2^WT^ and FOXL2^C134W^. Analysis of the resulting datasets revealed FOXL2^C134W^ binding to a majority of FOXL2^WT^ binding elements (80%) and, strikingly, to a unique set of genomic regions. Motif analysis of the uniquely bound regions showed an enrichment for SMAD3^17^ motifs at FOXL2^C134W^ bound sites as compared to FOXL2^WT^ and revealed that the genomic elements uniquely bound by FOXL2^C134W^ were enriched in a novel motif not observed in the elements common to FOXL2^WT^ and FOXL2^C134W^, supporting a model of altered DNA binding specificity. We confirmed FOXL2^C134W^ specificity to novel DNA binding elements *in vitro* and analysis of histone modifications associated with active enhancers confirmed that DNA elements uniquely bound by FOXL2^C134W^ adopted an active enhancer state following FOXL2^C134W^ induction. Integrated analysis of FOXL2^C134W^ specific proximal promoter binding sites and differential gene expression identified direct targets of FOXL2^C134W^ with therapeutic potential, including *SLC35F2*, a solute transporter whose expression has been previously shown to sensitize cells to the proapoptotic agent, YM155^18^. Treatment of our model with YM155 showed that FOXL2^C134W^ induction increased YM155 sensitivity relative to FOXL2^WT^. Our results thus provide a mechanistic understanding of FOXL2^C134W^ pathogenicity in AGCT and suggest its unique transcriptional targets may provide a route to the discovery of therapeutic vulnerabilities.

## Results

### Generation of isogenic FOXL2^C134W^ and FOXL2^WT^ cell lines

To address limitations of previous studies relying on overexpression of FOXL2^C134W^, we engineered tetracycline (TET)-on lentiviral constructs expressing FOXL2^WT^ or FOXL2^C134W^ under the control of doxycycline (Dox) (**Fig. 1A**). FOXL2^WT^ and FOXL2^C134W^ coding regions were fused to a V5 tag to distinguish exogenous from endogenous FOXL2. Stable cell lines were generated from the resulting viral constructs by transduction of the SV40 large T antigen immortalized human granulosa cell line (SVOG3e)^19,20^, followed by selection for viral integration. A concentration of Dox inducing the expression of FOXL2^C134W^ to the same level as observed by quantitative mass spectrometry in an AGCT cell line^21^ (**Fig. S1**) was identified (2.3:1, FOXL2^C134W^:FOXL2^WT^) (**Fig. 1B**). We named these cell lines as SVOG3e-WT, SVOG3e-MUT, and SVOG3e-EV for the empty vector control.

**Figure 1.**
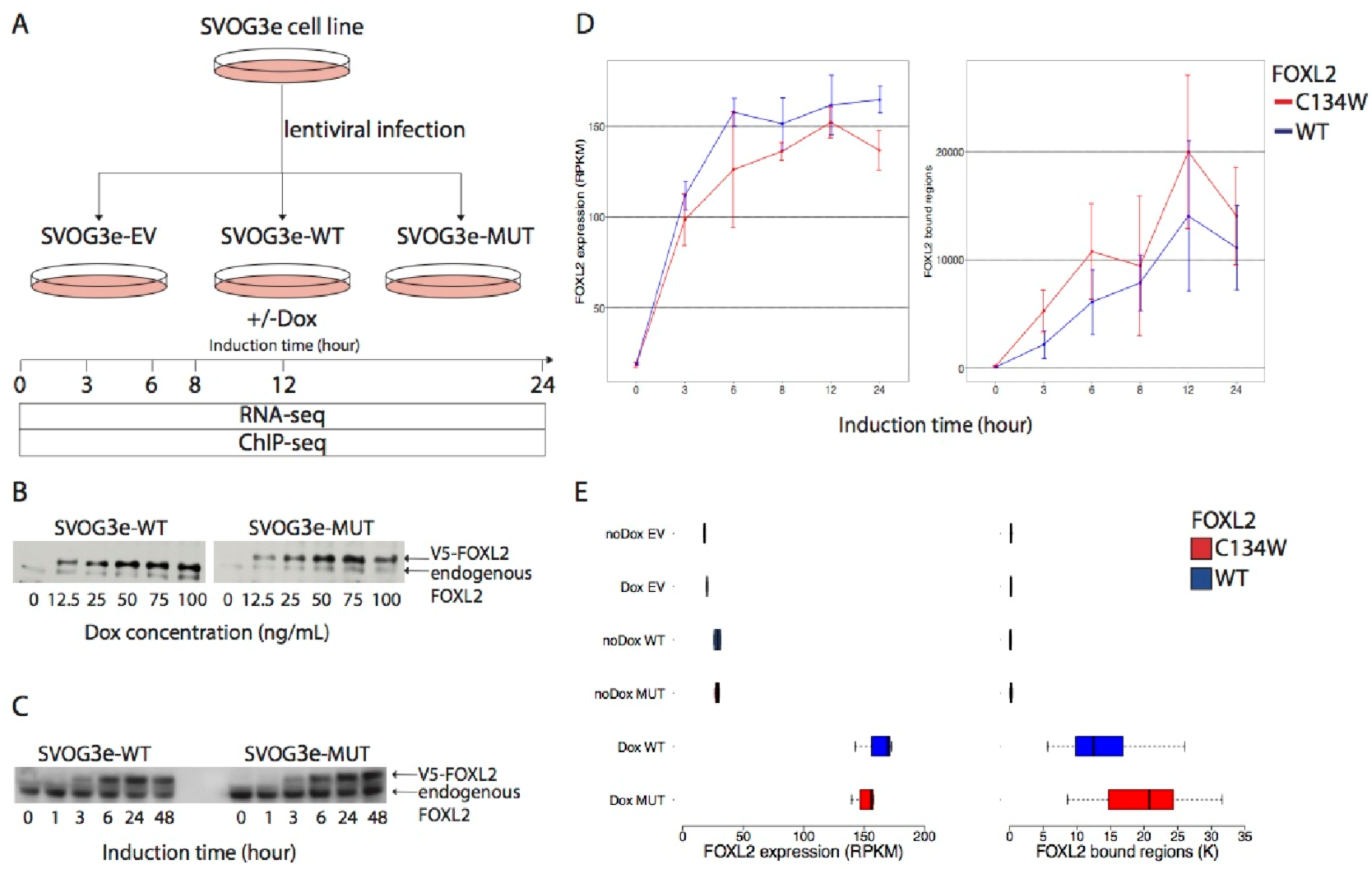
Generation of isogenic FOXL2^C134W^ and FOXL2^WT^ cell lines. **(A)** Schematic of the experimental design. SVOG3e-EV (empty vector), SVOG3e-WT (FOXL2^WT^), and SVOG3e-MUT (FOXL2^C134W^) are the three inducible cell lines used in this study. (**B**) Western blot of FOXL2^WT^ and FOXL2^C134W^ protein expression over Dox concentration. (**C**) Western blot of V5-FOXL2^WT^ and V5-FOXL2^C134W^ protein expression over Dox induction time. (**D**)Expression of *FOXL2* over time in SVOG3e-WT (blue) and SVOG3e -MUT (red) as assessed by RNA-seq (left panel) (**Fig. S2**). FOXL2 bound regions over time in SVOG3e -WT (blue) and SVOG3e -MUT (red) as assessed by V5-FOXL2 ChIP-seq (right panel). (**E**) *FOXL2* expression as assessed by RNA-seq (left panel) (Table S1) and genome-wide number of MACS2 enriched regions derived from V5-FOXL2 ChIPseq (right panel) at 12 hrs of Dox induction. The four control samples are non-induced and induced empty vector controls (‘noDox EV’, ‘Dox EV’), non-induced SVOG3e-WT (‘noDox WT’) and noninduced SVOG3e-MUT (‘noDox MUT’)). The two experimental samples are induced SVOG3eWT (‘Dox WT’) and induced SVOG3e-MUT (‘Dox MUT’). Left panel has the following number of replicates in ‘noDox EV’, ‘Dox EV’, ‘noDox WT’, ‘noDox MUT’, ‘Dox WT’ and ‘Dox MUT’: 2, 3, 2 (outlier is removed), 4, 3, 4 respectively. Right panel: 3, 3, 3, 4, 3, 4 replicates, respectively

To examine the dynamics of FOXL2^WT^ and FOXL2^C134W^ induction we synchronized SVOG3e-WT and SVOG3e-MUT cell lines and harvested cells at 0, 3, 6, 8, 12 and 24 hours (hrs) following Dox induction for RNA-seq and ChIP-seq analysis. The RNA-seq time series revealed a linear increase in the transcript levels of FOXL2^WT^ and FOXL2^C134W^ following Dox treatment until 6 hrs at which point expression remained constant until 24 hrs post induction (**Fig. 1D, Table S1**). Western blots of protein extracts from a matched time series using an anti-V5 antibody confirmed a similar expression pattern (**Fig. 1C**).

To examine the DNA binding dynamics of FOXL2^WT^ and FOXL2^C134W^ we performed ChIPseq using an anti-V5 antibody across the time series. Consistent with transcript and protein expression we observed an increase in the chromatin occupancy of FOXL2^WT^ and FOXL2^C134W^ over time with a maximum occupancy observed 12 hrs post-induction, 6 hrs past the maximum transcript expression (**Fig. 1D**). We selected 12 hrs post-induction for further comparisons and performed anti-V5 ChIP-seq and RNA-seq profiling at 0 hrs and 12 hrs following Dox induction in triplicates. Both FOXL2^WT^ and FOXL2^C134W^ showed a similar expression level 12 hrs postinduction (median RPKM 169.8 and 155.6, respectively) and a strong increase in DNA binding as compared to controls (**Fig. 1E)**. We confirmed that the level of expression of FOXL2^C134W^ mRNA in our models at 12 hrs is similar to that seen in primary AGCT^7^ (median RPKM 106.9) (**Fig. S3**, Table **S1**).

### FOXL2^C134W^ induction drives a unique transcriptional program

We next examined global transcription changes following induction of FOXL2^WT^ or FOXL2^C134W^. To control for the effects of Dox alone we compared expression in both cases to our vector control following 12hrs of Dox treatment. A similar number of up and down-regulated protein coding genes were observed upon FOXL2^WT^ or FOXL2^C134W^ induction consistent with its known dual role as both a transcriptional activator and repressor^22^ (**Fig. 2A** and **2B**). We confirmed known FOXL2 ^WT^ transcriptional targets including *STAR, CDKN2B*, and *INHBB* (Table **S2**). Interestingly a significant fraction (58.6%) of differentially expressed genes found following FOXL2^C134W^ induction did not overlap with those found following FOXL2^WT^ induction (**Fig. 2B**) suggesting that FOXL2^C134W^ drives a distinct transcriptional program.

**Figure 2.**
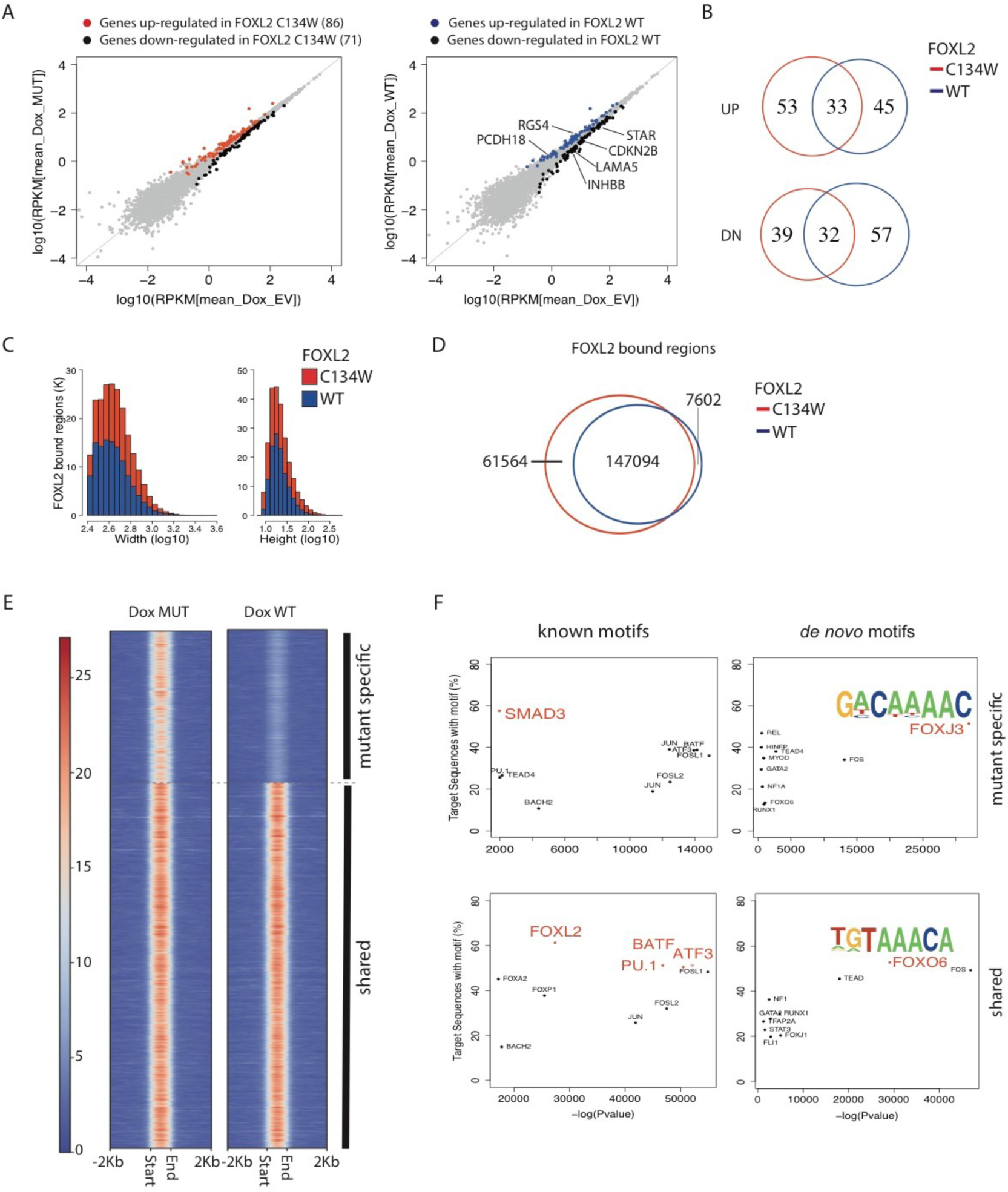
The gene expression profile and DNA binding specificity are altered by FOXL2^C134W^. (**A**) Scatter plots showing gene expression in Dox MUT versus Dox EV (left panel) and Dox WT versus Dox EV (right panel). Differentially expressed genes (FDR 0.05) are highlighted in red (genes up-regulated in Dox MUT), blue (genes up-regulated in Dox WT) and black (genes down-regulated). Examples of known FOXL2 targets are labeled. (**B**) Venn diagrams comparing the up-regulated (upper) and down-regulated (bottom) genes in Dox WT (blue) or Dox MUT (red) versus Dox EV. Only genes that are up- or down-regulated in minimum 3 out of the 4 Dox MUT replicates, and in minimum 2 out of the 3 Dox WT replicates were included (Table S2). (**C**) Histograms displaying width (left) and pileup height at peak summit (right) of MACS2 peak calls for V5-FOXL2 ChIP-seq in Dox WT (blue) and Dox MUT (red). (**D**) Venn diagram displaying the overlap between MACS2 enriched regions called in Dox MUT and Dox WT. Dox MUT enriched regions were defined as present in Dox condition and that do not overlap with enriched region from Dox EV control or Dox WT. Dox WT enriched regions were defined as present in Dox condition and that do not overlap with enriched region from Dox EV control or Dox MUT. We used a threshold of minimum 50% reciprocal overlap. (**E**) Heat maps of RPKM values associated in Dox MUT (left) and Dox WT (right) with statistically called FOXL2 enriched regions. Two kilobases on each side of the peaks are shown. Heat maps are ordered based on the specificity of the FOXL2 peaks: mutant-specific enriched regions, enriched regions shared between wildtype and mutant (minimum 50% overlap reciprocally between Dox WT and Dox MUT). Dox WT and Dox MUT enriched regions were defined as not overlapping with enriched regions found in Dox EV. One representative replicate is shown (**Fig. S5** for all replicates). (**F**) Scatter plots of the top10 enriched known (left) and *de novo* (right) motifs in the mutant-specific peaks (61,564) (top row) defined as not having any overlap with enriched regions found in Dox EV or in Dox WT and in the shared peaks (147,094) (bottom row). *De novo* motifs are labeled according to the best match to known available motif from databases. Motifs with a frequency >=50% are highlighted in red (full list of known motifs in **Table S3**)

### FOXL2^C134W^ binds unique genomic regions

Having established that FOXL2^WT^ and FOXL2^C134W^ induction drives distinct transcriptional programs in our model we sought to determine whether this reflected an altered genomic occupancy. Enriched regions (ERs) derived from ChIP-seq using an anti-V5 antibody following 12 hrs of FOXL2^WT^ or FOXL2^C134W^ induction showed similar genome-wide distributions in average widths and normalized aligned read densities (**Fig. 2C**). We also observed a similar genomic distribution for both FOXL2^WT^ and FOXL2^C134W^ of putative binding elements with an enrichment within gene bodies and promoters (59% of the enriched regions) (**Fig. S4**). In addition to binding to a majority of genomic regions occupied by FOXL2^WT^ (∼80%) FOXL2^C134W^ was also found to bind a large collection of unique genomic sites (61,564 regions genome-wide; **Fig. 2D**).

To further investigate FOXL2^C134W^ binding specificity we performed a motif analysis of enriched genomic regions^23^. To identify DNA motifs specifically enriched in FOXL2^C134W^ bound regions we compared the most significantly enriched motifs within the 61,564 genomic regions uniquely bound by FOXL2^C134W^ to those within the 147,094 regions bound by both FOXL2^WT^ and FOXL2^C134W^ (**Fig. 2D-E** and **S5**). Motifs specifically enriched in the shared regions included FOX family related motifs (FOXL2, FOXA2 and FOXP1) as expected (**Fig. 2F**). Within the FOXL2^C134W^ specific regions, TEAD4 and SMAD3 motifs were found to be unique within the top 10 enriched motifs (**Fig. 2F**). SMAD3 is a known partner of FOXL2^14,17,24-28^, and we confirmed that the SMAD3 motif was also enriched in the shared regions albeit at a lower frequency (53% vs 58%) and significance compared to FOXL2^C134W^ (Table S3).

Predicted *de novo* enriched motifs in regions bound by both FOXL2^WT^ and FOXL2^C134W^ included the consensus forkhead binding element^29-33^ 5’-[(G/A)(T/C)(A/C)AA(C/T)A]-3’ as the most frequent motif identified (53% of the ERs) (**Fig. 2F** bottom-right panel) consistent with a model where FOXL2^C134W^ and FOXL2^WT^ bind to a common DNA motif. *De novo* motif detection within the FOXL2^C134W^ specific regions revealed that the most frequent motif (found in 51% of FOXL2^C134W^ specific regions) shared features of the FOXJ3 motif (**Fig. 2F**). Interestingly the FOXJ3-like motif was not observed within the shared regions suggesting that it might represent a novel DNA element bound by FOXL2^C134W^. To assess the relative positions of the *de novo* motifs within the enriched regions we plotted the distribution of the most frequent motifs across the FOXL2^C134W^ specific and shared regions and confirmed they were enriched within the centre of the enriched regions (**Fig. S6**).

### FOXL2^C134W^ induction is associated with distinct regulatory states

We next sought to examine the consequences of FOXL2^C134W^ induction on the epigenetically defined regulatory landscape. For this we performed ChIP-seq for H3K27me3, H3K27ac, H3K4me3, H3K4me1 in SVOG3e cells harvested 12 hrs following Dox induction of either FOXL2^WT^, FOXL2^C134W^ or empty vector. The resulting datasets were normalized to their respective input control and enriched regions identified^34^. No directional gain or loss of regulatory states as defined by these histone modifications, either alone or in combination, were observed following induction of either FOXL2^WT^ or FOXL2^C134W^ (**Fig. 3A**). However, distinct sets of chromatin states were observed for both FOXL2^WT^ and FOXL2^C134W^ after Dox induction, supporting a specific FOXL2^C134W^ transcriptional program (**Fig. 3B**). We observed the expected relationships between gene expression and presence of the active (H3K4me3) or repressive (H3K27me3) marks in the promoter of genes (**Fig. 3C**). However, a majority of these epigenetic state changes were not associated with direct FOXL2 recruitment suggesting that they represent secondary effects.

**Figure 3.**
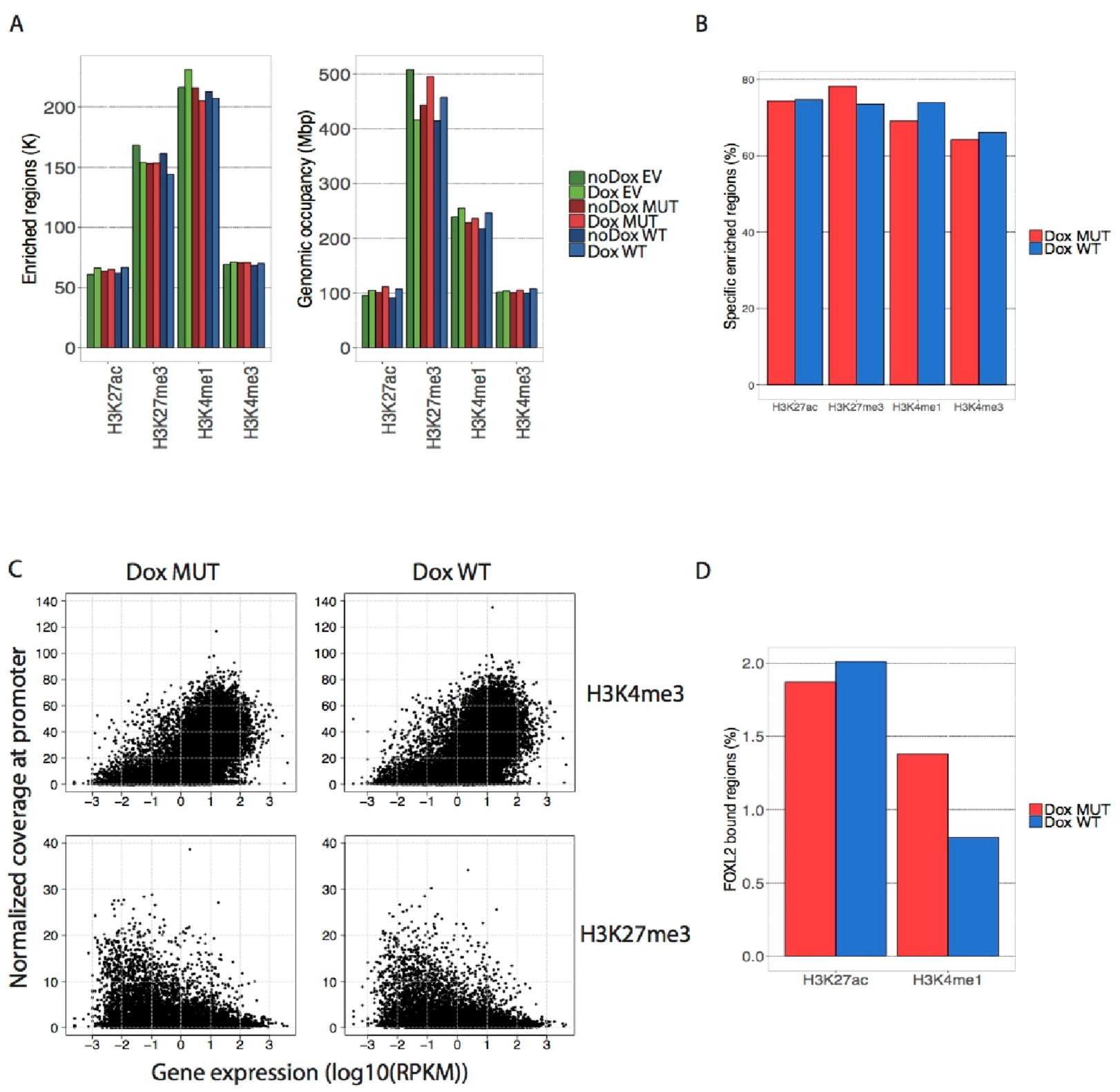
FOXL2^C134W^ induction is associated with distinct regulatory states. (**A**) Barplot displaying the number of ERs called (left panel) and the ER genomic occupancy (right panel) of each histone modification in the six different conditions. (**B**) Barplots showing the percent of ERs in Dox MUT that do not overlap with ER in Dox WT or vice versa for each examined histone modification. A genome-wide window-based approach (500 base pairs bins) was used. Only ERs that do not overlap with any ER from empty vector control were considered. (**C**) Scatterplots of H3K4me3 or H3K27me3 normalized coverage at promoter (transcription start site (TSS) +/-2Kb) versus gene expression (RPKM) in Dox WT and Dox MUT. (**D**) Barplots showing the percent of FOXL2 bound regions overlapping with ERs called for H3K4me1 and H3K27ac.

Epigenetic modifications including H3K4me1 and H3K27ac mark active enhancer states and thus we asked to what degree they were associated with the binding of FOXL2^C134W^. Surprisingly, only 1.9% of FOXL2^C134W^ binding events were associated with H3K27ac and 1.4% with H3K4me1 suggesting that a majority of the FOXL2^C134W^ binding did not result in the recruitment of histone methyl/acetyl-transferases and the establishment of an active enhancer state (**Fig. 3D**). This is consistent with a proposed model where a majority of FOXL2 binding events are transitory and non-functional^35^.

We then examined to what extent the genomic sites containing the mutant-specific DNA binding motif discovered *de novo* were occupied by FOXL2^C134W^. We searched genome-wide for the presence of this motif and only considered those overlapping with active chromatin states (defined as H3K27ac or/and H3K4me1) (5,867 out of 735,832). Only 5.7% of such motifs (335 out of 5,867) were bound by FOXL2^C134W^ suggesting that many potential DNA binding sites of FOXL2^C134W^ remain unbound, in accordance with previous studies on FOXA1 and FOXA2 transcription factors^36-39^.

### FOXL2^C134W^ binds a novel motif *in vitro*

Having established a DNA motif specific to FOXL2^C134W^ ChIP-seq enriched regions we sought to confirm whether FOXL2^C134W^ specifically bound the motif *in vitro*. Representative examples of mutant-specific motifs and shared motifs located in the promoter regions of *IL6* and *CHRNA9* (specific to FOXL2^C134W^) (**Fig. 4A** and **4B**) and *NFE2L2* (shared by FOXL2^WT^ and FOXL2^C134W^) (**Fig. 4C**) were identified and used to design electrophoretic mobility shift assay (EMSA) probes (**Fig. 4D**). Nuclear protein extracts of SVOG3e-WT and SVOG3e-MUT cell lines were prepared following Dox induction and incubated with IRDye 700 labelled probes (see methods). As predicted from the ChIP-seq datasets, we observed a mobility shift in the *IL6* and *CHRNA9* promoter DNA labeled probes incubated with FOXL2^C134W^ containing nuclear extracts but not in the probes incubated with the control (no extract) or FOXL2^WT^ induced extracts. In contrast, both FOXL2^WT^ and FOXL2^C134W^ nuclear extracts shifted the *NFE2L2* promoter DNA labeled probe consistent with the motif being bound by both FOXL2^WT^ and FOXL2^C134W^ *in vivo*. To confirm the observed binding was specific to the DNA motifs tested, we introduced competing unlabeled probes either identical to the labeled probe or containing a mutation in the conserved DNA motif. In the presence of unlabeled competing probe, the probe shift was eliminated in both the FOXL2^C134W^ specific and shared probes. In contrast the mutated probes had no effect (**Fig. 4D**).

**Figure 4.**
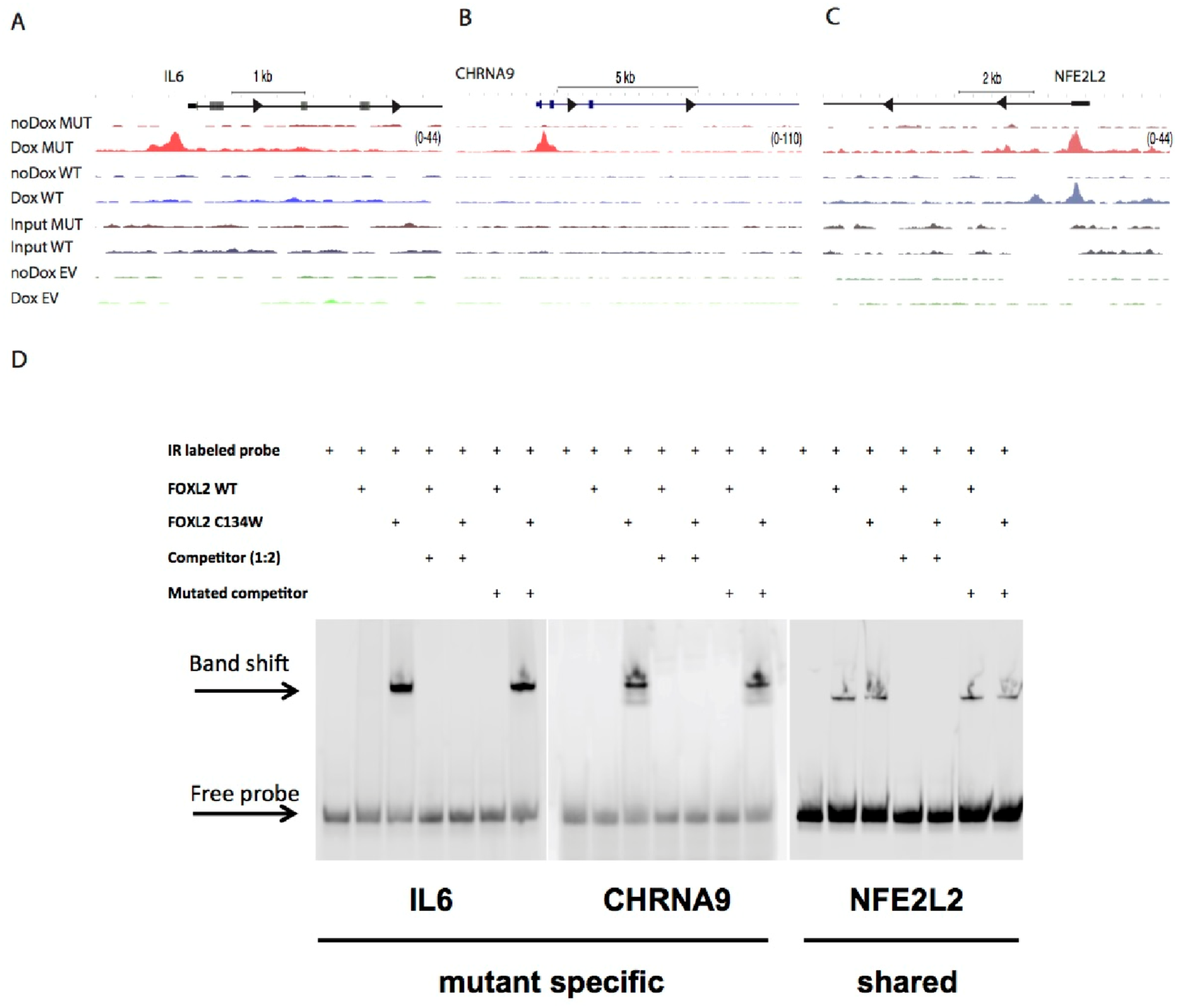
FOXL2^C134W^ binds specifically a novel motif *in vitro.* UCSC genome browser tracks highlighting the (**A**) *IL6* (chr7: 22,765,500-22,770,250), (**B**) *CHRNA9* (chr4: 40,334,53040,346,678), and (**C**) *NFE2L2* (chr2:178,121,500-178,130,750) gene promoters. One representative replicate is shown for each condition. (**D**) EMSA analysis of FOXL2^WT^ and FOXL2^C134W^ binding to *IL6, CHRNA9* and *NFE2L2* promoters. *CYP19A1* (aromatase PII promoter) was used as a positive control (**Fig. S7**)^12^.

### Identification of FOXL2^C134W^ direct target genes

Although we identified 61,564 uniquely bound DNA elements for FOXL2^C134W^, the vast majority of these were found to be not marked as active enhancers, defined as regions marked by H3K4me1 and H3K27ac, and were not associated with a directional transcriptional change when comparing FOXL2^WT^ to FOXL2^C134W^ arguing that the vast majority of FOXL2 binding events do not lead directly to transcriptional changes. To rule out the possibility that FOXL2^WT^ and FOXL2^C134W^ play a major role in regulating non protein-coding transcripts we analyzed RNA-seq libraries generated by ribodepletion from total RNA extracted from SVOG3e-WT, SVOG3e-MUT and SVOG3e-EV engineered cells harvested at 12 hours following Dox induction. FOXL2^WT^ induction resulted in a similar number of up- and down-regulated noncoding genes (52 and 49, respectively) consistent with patterns observed for protein-coding genes. FOXL2^C134W^ induction resulted in a similar split of down- and up-regulated genes (126 versus 93). Out of the 3,521 FOXL2^C134W^ uniquely bound DNA elements located within the proximity of noncoding genes (noncoding gene start +/- 2Kb), only ten were found to overlap differentially expressed non-coding genes. Two were up-regulated (*HIST3H2BA, SNORD27*) and eight were down-regulated (*SNORD22* and seven unannotated non-coding genes). Interestingly snoRNAs have been proposed to play an important role in the progression of cancer^40^. Taken together our analysis suggests that specific binding of FOXL2^C134W^ in our model has very little direct impact on both coding and non-coding gene expression.

However, given its pathognomonic relationship to AGCT we reasoned that the few direct targets identified in our model may be relevant to understanding AGCT pathology and reveal potential therapeutic vulnerabilities. To identify protein-coding genes that were direct target of FOXL2^C134W^ we identified genes with FOXL2^C134W^ specific binding events in their promoter regions and significantly up- or down-regulated following FOXL2^C134W^ induction, but not in any FOXL2^WT^ replicate or in the vector control. Of the FOXL2^C134W^ specific enriched regions overlapping with promoter regions (22% (13491/61564)) only four genes met our stringent criteria. Three genes were up-regulated (*CHRNA9, S1PR1* and *SLC35F2*) and one was down regulated (*ATP5E*). Strikingly, all four genes identified are highly relevant to cancer biology (**Table 1**). *CHRNA9* has been previously identified as highly expressed in breast cancer^41^; *S1PR1* is a G-protein coupled receptor involved in the cytokine signaling in the immune system, and is known to be up-regulated in ovarian cancer tissues and cell lines^42^; *SLC35F2* is a potential oncogene highly expressed in non-small-cell lung carcinoma^43^, indispensable for papillary thyroid carcinoma progression^44^, and regulated by androgen receptor axis signaling in prostate cancer^45^; *ATP5E* is known to be down regulated in papillary thyroid cancer^46^. These findings prompted us to examine whether those genes were also expressed in primary AGCTs^7^. Three out of the four genes were expressed at a significant level (median RPKM across the four AGCT samples are: 12.96 (*S1PR1*), 9.01 (*SLC35F2*), 19.24 (*APT5E*)) suggesting that they may represent *bona fide* targets of FOXL2^C134W^ in a primary tumour setting.

**Table 1.** FOXL2^C134W^ direct target genes. List of coding genes containing a mutant–only ER in their promoter region as indicated (−1kb/+300bp, −2kb/+2kb or −5kb/+5kb upstream and downstream TSS) and are differentially expressed (FDR 0.05) in Dox MUT as compared to Dox EV but not differentially expressed in any Dox WT replicate. A minimum of three replicates was used to call differentially expressed genes and FOXL2 ERs.

|  | Gene | -log10 (FDR) (RNA) | Fold-change (RNA) | -log10 (q-value) (MACS2) | TSS region | Gene function | Relevance to cancer |
| --- | --- | --- | --- | --- | --- | --- | --- |
| Up | CHRNA9 | 4.15 | 7.14 | 28.30 | -1kb/+300bp | Ionotropic receptor | Hsieh et al 2014 |
| Up | S1PR1 | 5.02 | 1.72 | 18.07 | -2kb/+2kb | G-protein coupled receptor | Wen et al., 2015 |
| Up | SLC35F2 | 3.56 | 1.52 | 15.20 | -1kb/+300bp | Transmembrane drug/metabolite transporter | He et al 2018; Nyquist et al 2017 |
| Down | ATP5E | 4.08 | -1.62 | 21.84 | -5kb/+5kb | ATP synthase | Hurtado-Lopez et al, 2015 |

### FOXL2^C134W^ target provides a potential therapeutic vulnerability in AGCT

FOXL2^C134W^ induced targets are candidates for therapeutic intervention as they would be expected to show tumour specificity. *SLC35F2*, a putative direct target of FOXL2^C134W^, encodes for a solute transporter previously shown to transport sepantronium bromide (YM155), a transcriptional suppressor of survivin expression having activity against a broad range of cancer types^47,48^ (**Fig. 5A**). Our analysis suggests that FOXL2^C134W^ drives increased expression of *SLC35F2* and thus we hypothesized that tumour cells expressing FOXL2^C134W^ would show increased sensitivity to YM155. To test this hypothesis, we induced SVOG3e-WT, SVOG3e-MUT and SVOG3e-EV with Dox for 12 hrs and treated the cells with increasing concentrations of YM155 and measured cell viability over time. This analysis revealed an increased sensitivity to YM155 for SVOG3e cells expressing FOXL2^C134W^ compared to cells expressing FOXL2^WT^ or the vector control (**Fig. 5B**). This effect was significant (one-way ANOVA) at both 15 and 39 hrs at 20nM (*P*-value 0.001) supporting our hypothesis that FOXL2^C134W^ drives increased *SLC35F2* expression leading to an increased sensitivity to the YM155. Collectively, these results suggest YM155 may be an effective therapeutic strategy for AGCT (**Fig. 5C**).

**Figure 5.**
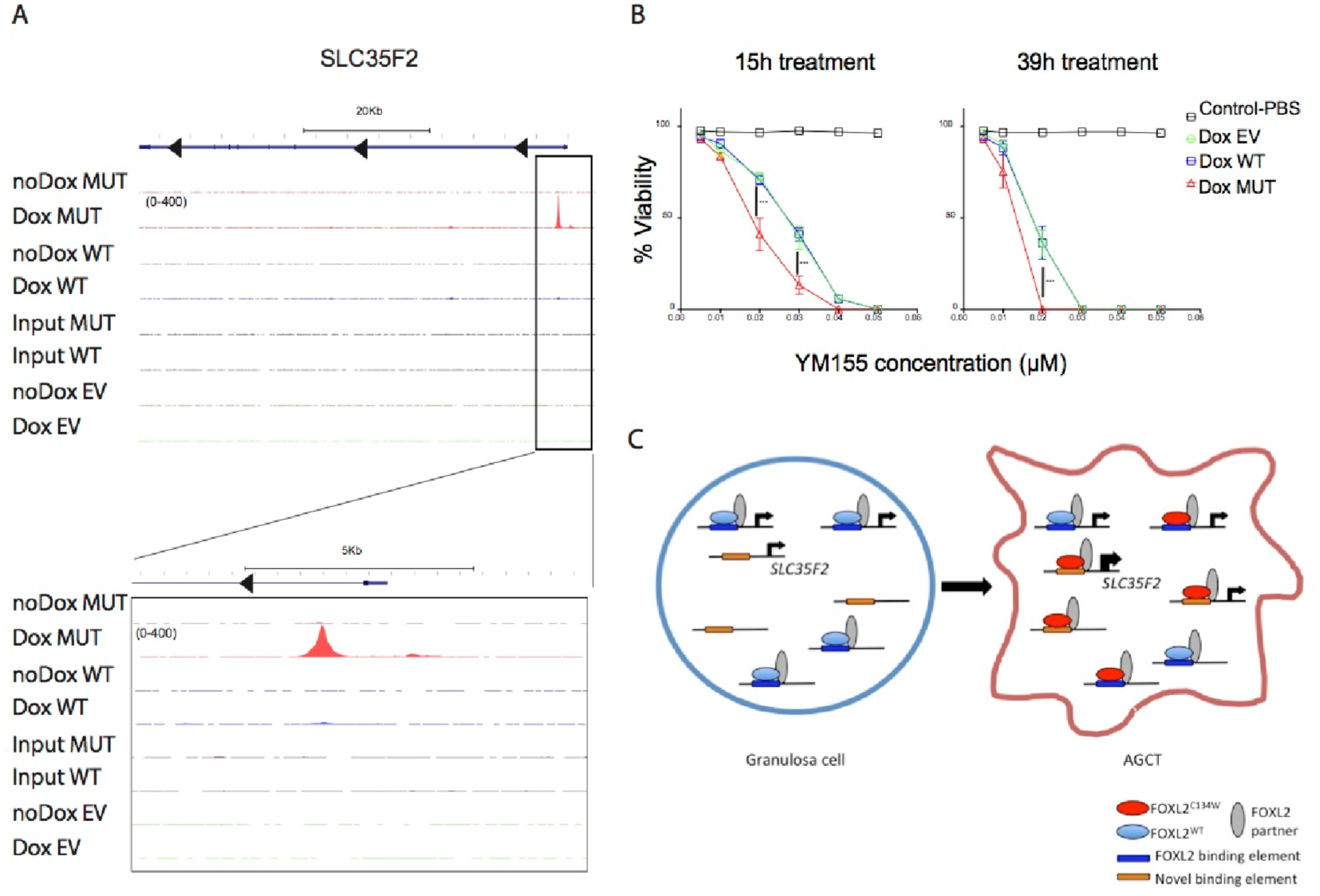
FOXL2^C134W^ target provides a potential therapeutic vulnerability in AGCT. (**A**) UCSC genome browser screenshot showing FOXL2 ChIP-seq tracks at *SLC35F2* locus (chr11:107,661,717-107,729,914). One representative replicate is shown for each condition. Below panel is a zoom in *SLC35F2* promoter region (TSS+/-5Kb). (**B**) Cellular viability of Dox WT, Dox MUT and Dox EV after 15h and 39h of Ym155 treatment with various concentrations of YM155 (10, 20, 30, 40, 50 nM) – after 12 hrs Dox induction. Experiments were performed in triplicate. Data are represented as mean ± standard deviation (**C**) Model of FOXL2^C134W^ mechanism in AGCT pathogenesis.

## Discussion

The somatic missense point mutation c.402C>G (p.C134W) in FOXL2 is pathognomonic for AGCT^7^. Understanding the molecular consequences of FOXL2^C134W^ expression has important therapeutic implications in AGCT. In this study, we undertook to measure the FOXL2^C134W^ expression in a controlled model to elucidate the overall molecular signature and identify direct transcriptional targets. For this we developed a series of model cell lines engineered with inducible V5 tagged FOXL2^WT^ and FOXL2^C134W^ and performed comprehensive ChIP-seq and RNA-seq analysis over an induction time series. Analysis of the resulting datasets showed that FOXL2^C134W^ was bound to a majority of FOXL2^WT^ elements and was associated to over 60k novel regions genome wide. We confirmed previous reports suggesting that the vast majority of FOXL2 binding events (>95%) could not be associated with transcriptional changes supporting the hypothesis that these represent examples of “non-functional” or transient transcription factor recruitment events^35^. Examination of the epigenetic state of chromatin flanking FOXL2^C134W^ recruitment sites further supported this hypothesis demonstrating that a majority of sites are not epigenetically marked as active enhancer states. Chromatin elements found to be epigenetically modified following FOXL2^C134W^ recruitment did not show enrichment in association with differentially expressed genes nor a significantly different genomic annotation.

*De novo* motif analysis revealed that FOXL2^C134W^-specific binding sites are enriched in a novel DNA motif and this motif was shown to be specifically bound by FOXL2^C134W^ *in vitro*. Comparison of bound regions to known motifs suggests FOXL2^C134W^ may also have an enhanced interaction with SMAD3 as compared to FOXL2^WT^, in agreement with what has been previously suggested by structural modeling^6,7^ and recently validated by co-IP^17^ (Belli et al. 2018). Overall, our results suggest that FOXL2^C134W^ contributes to a pathogenic transcriptional signature through recruitment to novel DNA elements genome-wide - a subset of which alter the expression of genes supporting cellular transformation.

Integration of RNA-seq and ChIP-seq datasets following FOXL2^C134W^ induction allowed us to predict with high confidence four direct target genes. While our methods of gene assignment were conservative, such a low number of direct gene targets demonstrating differential expression is somewhat unexpected given the central role of FOXL2 ^C134W^ in AGCT. Indeed, we were unable to discern any specific pathways from the small number of differentially expressed genes following FOXL2^C134W^ induction. This could be in part driven by an inability of the SVOG3e cell line to respond appropriately to FOXL2^C134W^, or this may indeed represent a full repertoire of FOXL2^C134W^ targets. Regardless, the small number of putative targets identified have all been implicated as key drivers of transformation and thus are likely to be of relevance in adult granulosa cell tumorigenesis. Importantly we were able to validate *SLC35F2* as a direct target of FOXL2^C134W^ through an increase in sensitivity to YM155, a compound suppressing survivin and currently the subject of 11 clinical trials (clinicaltrials.gov). Our study provides a molecular mechanism for YM155 sensitivity driven by FOXL2^C134W^ and thus provides a foundation for additional studies designed to test the effectiveness of YM155 as therapeutic strategy in the management of adult granulosa cell tumors.

## Acknowledgements

This work was supported by the Terry Fox Research Institute Program Project Grant (#1021) to S.S., G.B.M., D.G.H. and M.H. This research was enabled in part by support provided by WestGrid and Compute Canada (www.computecanada.ca) and Canada Foundation of Innovation (#31343 & #31098). The authors also wish to acknowledge Canada’s Michael Smith Genome Sciences Centre, Vancouver, Canada for computational resources and support. A full list of funders of infrastructure and research supporting the services accessed is available at www.bcgsc.ca/about/funding_support.

## Author Contributions

Conceptualization: G.B.M, D.G.H., and M.H.; Methodology: G.T-G, S.-W.G.C and G.M; Investigation, G.T-G., S.-W.G.C., Q.C. and M.M.; Software: A.C. and M.B.; Formal analysis, A.C and M.H; Supervision: M.H; Writing: original draft, A.C. and M.H.; Writing: review & editing, A.C., M.H., S.-W.G.C and G.M..

## Declaration of Interests

The authors declare no competing financial interests.

## Data Accession

Sequence datasets generated as part of this study have been deposited to GEO under the accession GSE126171.

## Methods

### Cell lines and cell culture

A non-tumorigenic immortalized human granulosa-lutein cell line (SVOG3e) was generated by transfecting human primary granulosa-lutein cells with plasmids expressing the simian virus 40 large T antigen and hTERT^19,20^. SVOG3e cells were grown in media 105/199 supplemented with 5% defined fetal bovine serum (Hyclone SH 30070.03). Media 105/199 was prepared mixing equal volumes of media MCDB 105 (Sigma M6395) and 199 (Sigma M5017) which were prepared according to manufacturer’s protocol. HEK293T cells were cultured in DMEM (Gibco 11995-065) supplemented with 10%FBS (Gibco 12484-028).

### DNA Constructs

FOXL2^WT^ cDNA was kindly provided by Dr Margareta Pisarska (UCLA). QuickChange II sitedirected mutagenesis (Agilent) was used to introduce a single mutation C402G to obtain a FOXL2^C134W^ mutant. Following sequencing, PCR was performed on the FOXL2^WT^ and FOXL2^C134W^ constructs to generate pDONR221/wtFOXL2 and pDONR221/FOXL2/C134W entry vectors, using the Gateway BP Clonase II Enzyme Mix (Invitrogen 11789-020). Genes were transferred into the lentiviral vector pLVX-Tight-Puro (Clontech) modified to contain a C-terminal V5 tag (pLVX-V5, kindly provided by Dr. S. Aparicio) using the Gateway LR Clonase II Enzyme Mix (Invitrogen 11791-020).

### Lentivirus and production of stable cell lines

To generate tetracycline inducible FOXL2 cell lines, we used the Lenti-X TET-ON Advanced Inducible Expression System (Clontech), which would allow controlled expression of FOXL2^WT^ or FOXL2^C134W^ protein by addition of Dox (Sigma, D9891). Replication-defective lentivirus containing the rtTA Advanced TET-ON was generated by co-transfection of pLVX-TET-ON, pCMVΔR8.91 and pMD2VSV-G envelope plasmids into HEK293T cells using TransIT-LTI transfection reagent (Mirus Bio). Virus particles collected at 48 hours post-transfection were passed through a 0.45 μm filter and added to SVOG3e cells at a M.O.I. of 3. Cells were selected with 0.6 mg/ml Geneticin (Gibco 10131-035) over 2 weeks, generating a polyclonal SVOG3e/rtTA cell line. pLVX-V5, pLVXV5-wtFOXL2 and pLVX-V5-FOXL2/C134W lentiviruses were generated in the same manner as described above, except that each virus was added to SVOG3e/rtTA cells and selected with 0.5 μg/ml Puromycin (Sigma P8833) and 0.6 mg/ml Geneticin to create polyclonal stable cell lines SVOG3e/rtTA/V5 (V5 control cell line), SVOG3e/rtTA/wtFOXL2 (inducible Wild Type FOXL2 cell line) and SVOG3e/rtTA/FOXL2/C134W (inducible mutant FOXL2/C134W cell line).

### Cell cycle synchronization and inducible expression of FOXL2

Cell cycle synchronization was achieved by using the Double Thymidine Block method^49^. Briefly, SVOG3e/rtTA/V5, SVOG3e /rtTA/FOXL2^WT^ or SVOG3e /rtTA/FOXL2^C134W^ cells were seeded in SVOG3e media without antibiotics. 48 hours after seeding the cells, media was exchanged for media containing 2mM Thymidine and cells were incubated at 37C for 19 hours. To release the first block, media was removed and cells were washed three times with distilled Phosphate Buffered Saline (dPBS). Fresh media containing no Thymidine was added to the cells and the cells were incubated for another 9 hours. 2mM Thymidine was then added and the cells were cultured for additional 16 hours. After the cells were released from the 2nd block by removing the thymidine containing media and washing the cells with dPBS, Fluorescence Activated Cell Sorting (FACS) analysis was performed at various time points after the cells were released from the second cell cycle block and it was established that cells remained synchronized for up to 8 hours. By 24 hours, the cells were asynchronous.

Prior to Doxycycline induction, synchronized cells were washed twice with dPBS before induction of FOXL2 protein expression by the addition of media containing 12.5 ng/ml Doxycyclin. This concentration of Doxycycline was tailored such that the expression of ectopic V5-FOXL2 would be similar to endogenous FOXL2 expression levels in SVOG3e cells, as assessed by Western blot using a polyclonal goat anti-FOXL2 antibody (Novusbio NB100-1277), followed by ECL detection using anti-goat IgG-HRP (Santa Cruz sc-2350) (**Fig. 1B** and **Fig. S1**). Protein expression was detected as early as 3 hours post-induction.

### Cell line induction and collection of samples for RNA-seq and ChIP-seq

To monitor the effect of FOXL2^WT^ and FOXL2^C134W^ expression on DNA binding and protein expression over time, a time-course experiment was performed to look at FOXL2^WT^ or FOXL2^C134W^ binding on DNA by Chromatin-Immunoprecipitation followed by DNA sequencing (ChIP-seq). SVOG3e/rtTA/V5-wtFOXL2 or SVOG3e/rtTA/V5-FOXL2/C134W were seeded at 1×106 cells per 15 cm dish. Double Thymidine block was started 48 hours later after seeding the cells. After release from the second Thymidine block (T=0), media containing 12.5 ng/ml doxycycline was added. The induction with doxycycline was stopped at time points 3, 6, 8, 12 and 24 hours by removing the media by aspiration and washing the cells with cold dPBS. Non-induced control cells were harvested at T=0 and T=12 hours. Approximately 30 million cells were collected per time point. 5 million cells per time point were harvested for RNA preparation, protein assay and for western blot analysis. 25 million cells per time point was crosslinked with 1% formaldehyde (Sigma F8775) for 10 minutes at room temperature. Crosslinked cells were quenched with onetenth volume of 1.25 M Glycine (Sigma G7126) for 5 minutes and washed twice with dPBS before harvesting the cells. Cells were pelleted at 3,000 rpm, the supernatant removed and the cell pellet frozen at −80C. Additionally the control cell line SVOG3e/rtTA/V5 not expressing V5-FOXL2 was seeded and treated as above for one time point at 12 hours for non-induced and doxycyclineinduced cells. We used 3 replicates of FOXL2^WT^, 3 replicates of empty vector control and 4 replicates of FOXL2^C134W^ for all non-induced and induced samples across all time points for both RNA-seq and ChIP-seq, with the exception of the 3 hours induced FOXL2^C134W^ RNA-seq samples done in 3 replicates and the 12 hours non-induced empty vector control RNA-seq done in 2 replicates.

### Multiple reaction monitoring (MRM) analysis of induced FOXL2 expression

FOXL2^WT^ and FOXL2^C134W^ protein expression were quantified by targeted mass spectrometry. Isotopically labeled heavy peptide standards were used to generate a standard curve as previously described (Yap DB Blood 2011). FOXL2^WT^ and FOXL2^C134W^ proteins were isolated from KGN cells by immunoprecipitation with anti-FOXL2 antibodies and the amount of wild-type, mutant and common peptides was determined using the targeted mass spectrometry assay. We determined that the amount of FOXL2^C134W^ peptide was roughly 2 fold higher than the wild-type peptide.

### RNA-sequencing

Cells were harvested in dPBS by scraping, transferred to an Eppendorf tube and spun down at 900 x g for 5 minutes. The supernatant was discarded and the cell pellet resuspended in Qiazol Lysis reagent (Qiagen 79306). Total RNA was isolated using RNeasy kit (Qiagen 74106) as per the manufacturer’s protocol.

PolyA RNA-seq library construction was performed according to standard operating procedures available at http://www.epigenomes.ca/protocols-and-standards. For ribodepleted RNA-seq library construction, see next section. All libraries were sequenced on Illumina HiSeq 2500 (paired-end 75nt).

RNA-seq paired-end reads were aligned using BWA-backtrack v0.5.7 (polyA RNA-seq) or BWA mem v0.7.6 with options ‘-M’ and ‘-P’ (ribodepleted RNA-seq) and default parameters (Li and Durbin 2010) to a transcriptome reference consisting of the reference genome (GRCh37-lite) extended by read-length-specific exon–exon junction sequences. The sorted bam files were repositioned using JAGuaR^50^ v1.7.5 to a custom transcriptome reference (built with GRCh37-lite and Ensembl v69 (GenCode v14)) transforming reads spanning exon–exon junctions into largegapped alignments. The number of mapped reads produced was between 11 x 10^6^ and 59 x 10^6^ across poly-A RNA-seq libraries and between 88 x 10^6^ and 98 x 10^6^ across ribodepleted RNAseq libraries.

Using repositioned reads, we generated genome-wide coverage profiles (wiggled files) with a custom BAM2WIG java program (http://www.epigenomes.ca/tools-and-software) for further analysis and visualization in genome browser^51^. To generate those profiles, we included pairs marked as duplicated as well as pairs mapped in multiple genomic locations.

A custom RNA-seq quality control and analysis pipeline was applied to the generated profiles and a number QC metrics were calculated to assess the quality of RNA-seq library such as intron– exon ratio, intergenic reads fraction, strand specificity, 3’-5’ bias, GC bias and reads per kilobase of transcript per million reads mapped (RPKM) (Table S1). To quantify the exon and gene expression we calculated RPKM (Mortazavi A. *et al*. Nature methods 2008) metrics. For the normalization factor in RPKM calculations we used the total number of reads aligned into coding exons and excluded reads from mitochondrial genome as well as reads falling into genes coding for ribosomal proteins as well as reads falling into top 0.5% expressed exons. RPKM for a gene was calculated using the total number of reads aligned into its all merged exons normalized by total exonic length. The resulting files contain RPKM values for all annotated exons and coding and noncoding genes, as well as introns.

*FOXL2* expression before and after induction was used as quality control. One outlier behaving like an induced sample although labeled as non-induced has been removed. All possible pairwise comparisons between induced wildtype or mutant FOXL2 and induced vector alone after 12 hours Dox treatment were performed to identify differentially expressed genes using a custom DEfine v0.9.3 matlab tool (FDR 0.05, minimum RPKM 0.01, minimum number of reads per genes 25). For induced FOXL2^WT^ which had 3 replicates, we set a threshold of at least 2 out of 3 having at least one pairwise comparison that passes the FDR cutoff. For induced FOXL2^C134W^ which had 4 replicates, the threshold was 3 out of 4.

### Ribodepleted RNA-sequencing

RNA was extracted using a combination of mirVana miRNA Isolation kit (Thermofisher, AM1560) and All prep DNA/RNA Mini Kit (Qiagen, 80204). RNA was then assessed for quality and quantified using Agilent Bioanalyzer (Life Technologies). 500ng of total RNA was ribosomal RNA (rRNA) depleted using a NEBNext rRNA Depletion Kit (New England BioLabs, E6310L). First strand cDNA was generated using a Maxima H minus First Strand cDNA Synthesis Kit (Thermo Scientific, K1652) with the addition of 1μg of Actinomycin D (Sigma, A9415). The product was purified using in-house prepared 20% PEG in 1M NaCL Sera-Mag bead solution at a 1.8X ratio and then eluted in 35μL of Qiagen EB buffer. Second Strand cDNA was synthesized in a 50μL volume using SuperScript Choice System for cDNA Synthesis (Life Technologies, 18090-019) with 12.5mM GeneAmp dNTP Blend with dUTP. Double-stranded cDNA was purified with 20% PEG in 1M NaCL Sera-Mag bead solution at a 1.8X ratio and eluted in 40μL of Qiagen EB buffer, and fragmented using Covaris E220 (55 seconds, 20% duty factor, 200 cycles per burst). Sheared cDNA was end repaired/phosphorylated, single A-tailed, and adapter ligated using custom reagent formulations (New England BioLabs, E6000B-10) and in-house prepared Illumina forked small adapter. PEG (20%) in 1 M NaCl Sera-Mag bead solution was used to purify the template between each of the enzymatic steps. To complete the process of generating strand directionality, adapter-ligated template was digested with 5 U of AmpErase Uracil N-Glycosylase (Life Technologies, N8080096). Libraries were then PCR amplified and indexed using Phusion Hot Start II High Fidelity Polymerase (Thermo Scientific, F 549-L). An equal molar pool of each library was sequenced.

### FOXL2 ChIP-seq

Cell lysis, chromatin preparation, immunoprecipitation using anti-V5 antibody (Invitrogen, R960CUS) and recovery was performed as described previously^52^, with the following modifications; the chromatin pellet was resuspended in 400 μl of ChIP IP buffer and sonicated (Duty Cycle 20%, Intensity 8, Cycles/burst 200, Time: 60 seconds) in a Covaris Cov-3 Sonicator. Pre-cleared chromatin (150 μg) was incubated with 4 μl (2 μg) of monoclonal mouse anti-V5 antibody (Invitrogen R960-25) for 1 hr at 4C with rocking, followed by the addition of 40 μl of Protein G Dynabeads (Invitrogen) blocked with salmon sperm DNA and bovine serum albumin as previously described. Chromatin/antibody/Dynabead mixture was incubated overnight at 4C on a nutator. The Dynabead bound proteins and associated chromatin was washed and eluted as previously described and DNA was submitted to the GSC library core for library construction and sequencing. ChIP-seq library construction was performed according to standard operating procedures available at http://www.epigenomes.ca/protocols-and-standards.

Raw sequences were examined for quality, sample swap and reagent contamination using custom in house scripts. Sequence reads were aligned to NCBI GRCh37-lite reference using BWA-backtrack ^53^ v0.5.7and default parameters. The number of mapped reads was around 25 million. Custom java program (BAM2WIG) was used to generate wig files for downstream analysis and visualization^51^. Reads with BWA mapping quality scores <5 were discarded and reads aligned to the same genomic coordinate were counted only once in the profile generation. Enriched regions were called using MACS^34^ v2.1.1 (FDR 0.01) and default parameters. Input samples were used as control background. For normalized pileup height at peak summit, the option ‘—SPMR’ was used. Only ERs mapped to autosomes and sex chromosomes were included in the analysis. ERs having the same start and end positions were counted only once. FOXL2 ERs are defined as the ones called in Dox condition and that do not overlap with ER from Dox EV control. The overlap between genomic regions was generated using the intersect program of Bedtools^54^ v2.17.0. We used a threshold of minimum 50% of FOXL2 ER in the overlap (Fig 3D). Normalized coverage at promoter regions was computed using the wig files and a custom Java script.

Motif analysis (parameters for *de novo* motif discovery: size=given and length=8), motif occurrence plots (parameters: size 500, hist 5) and finding motif instances across the whole genome were done using Homer^23^ v4.8.2 findMotifsGenome, annotatePeaks and scanMotifGenomeWide Perl programs, respectively.

Read density heatmaps were generated using Deeptools bamCoverage, computeMatrix and plotHeatmap functions^55^. Plots were generated using R v3.3.2. Genome browser screenshots were downloaded from UCSC genome browser^56^ (http://genome.ucsc.edu).

### Histone modification ChIP-seq

Cells were trypsinized and collected by spin at 300g for 5 min and snap-frozen. Nucleosome density ChIP-seq^57,58^ was performed as described at https://www.jove.com/video/56085/generation-native-chromatin-immunoprecipitationsequencing-libraries. Libraries were sequenced on Illumina HiSeq 2500 (paired-end 125nt). Read sequences were aligned to NCBI Build 37 (hg19) human reference genome using BWA^53^ mem v0.7.6 with option ‘-M’. Enriched regions were called using MACS^34^ v2.1.1 ‘callpeak’ with FDR cutoff 0.01, ‘BAMPE’ and Grch37/hg19 effective genome size ‘2864785223’. Input samples were used as control background. Histone marks ERs are defined as the ones called in Dox condition and that do not overlap with ER from Dox EV control.

### Electrophoretic Mobility Shift Assay (EMSA)

Probes were designed using a 40bp sequence centered on the Homer derived motif in the ChIPseq derived FOXL2^WT^ and FOXL2^C134W^ shared peaks or FOXL2^C134W^ specific peaks located at proximal promoters. Probe sequences were shortened in case there were four consecutive G as per IDT manufacturer recommendation. Both sense and antisense DNA oligos were purchased and then annealed together to form a double-stranded DNA fragment. Probe sequences were double stranded and end Infra-Red labeled except the unlabeled probes. IRDye 700-labeled probes were designed so that they are centered on the motif (*IL6*: 5’-TTAGA**GTCTCAAC**CC-3’ *CHRNA9*: 5’-GGGCAGGACAGGGAA**GACAAAAC**AGATGATTCAACCGCA-3’ *NFE2L2*: 5’TAGCAATTGCGCAACAG**ATCAACA**GCTCCAACCTGTCCCT-3’). For competition assay, unlabeled unmutated probes and unlabeled mutated probes have been used (*IL6*: 5’-TTAGA**GTCTC<u>ggt</u>**CC- `3’`. *CHRNA9* 5’-GGGCAGGACAGGGAA**GAC<u>ggag</u>C**AGATGATTCAACCGCA-3’. *NFE2L2* 5’TAGCAATTGCGCAACAG**ATC<u>ggg</u>A**GCTCCAACCTGTCCCT-3’). All EMSAs were performed from the same nuclear protein extract and using the same binding conditions.

Cells were harvested 16 hours after Dox induction. Cells (5×10^6^) were treated with 200 μl of lysis buffer (50 mM KCl, 0.5% Nonidet P-40, 25 mM HEPES, pH 7.8, 2× proteinase inhibitor cocktail, 100 mM dithiothreitol) on ice for 5 min. After 1 min of centrifugation at 10,000 g, the supernatant was saved as a cytoplasmic extract. The nuclei were washed once with the same volume of buffer without Nonidet P-40 and then were put into a 100 ul of extraction buffer (500 mM KCl and 10% glycerol with the same concentrations of HEPES, proteinase inhibitor cocktail and dithiothreitol as in the lysis buffer) and pipetted several times. After centrifugation at 15,000 g for 5 min, the supernatant was harvested as the nuclear protein extract and stored at −80C. The protein concentration was determined with the bicinchoninic acid protein assay reagent (Fisher). IRDye 700 labeled oligonucleotide (LI-COR Biosciences) corresponding to specific consensus sequence was used. The binding reaction was performed using Odyssey Infrared electro-mobility shift assay kit (LI-COR Biosciences) according to the manufacturer protocol. Briefly, 10 μg of total nuclear protein was mixed with the designed oligonucleotide probe and left to bind for 30 minutes at room temperature in the dark. The binding reaction contains 1× binding buffer, 2mM DTT/0.25% Tween20, 0.05 μg Poly (dI-dC), 2.5% glycerol, 20 mM EDTA, 10 μg nuclear protein, 2.5 mM probe.

Protein-DNA complexes were resolved by electrophoresis on 5% polyacrylamide Tris/Acetate/EDTA (TAE) gel containing 0.5× TAE in the dark. Labeled complexes were detected according to the manufacturer’s protocol for the Odyssey CLx Infrared Imaging System (LI-COR Biosciences, Lincoln, NE). The ratio between dye-probe and no-dye-compete probe was 1:2 (10 uM : 20 uM). Binding incubation was for 55 minutes. Gel running time was 1.5 hr, 5% TAE gel in 0.5X TAE.

### Cell viability assay

SVOG3e/rtTA/V5Foxl2 cells were cultured in a 1:1 mix of media 199 (Sigma M5017) and media MCDB 105 (Sigma M6395) containing 5% fetal bovine serum (Hyclone Catalogue SH 30070.03). Cells were maintained in complete medium with 0.5 ug/ml Puromycin (Sigma P8833) and 0.6 mg/ml G418 (ThermoFisher, 10131035). Cells were seeded at 1.8 × 10^5^ / well in 6-well plate without Puromycin or G418 for 24 hours before induction with 12.5 ng/ml of doxycyclin for 12 hours. Cells were then treated with YM155 (Selleckchem, S1130, 10 mg) at 10 ∼ 50 nM concentration for 15 and 39 hours. Cells were collected and the cellular viability was tested using trypan Blue staining and counted with Countess II machine (LifeTechnologies). In order to assess the statistical significance across the three groups of cells (FOXL2^WT^, FOXL2^C134W^ and empty vector control), one-way Anova test was performed for every YM155 concentration (10, 20, 30, 40 and 50 nM) at both 15 and 39 hours of YM155 treatment. The test passed the threshold of *P*-value<0.001 at concentrations 20 nM and 30 nM after 15 hours of treatment and at concentration 20 nM after 39 hours of treatment.

## Supplementary Figures Legends

**Figure S1 related to Figure 1B.** FOXL2 standard curves for the (C134W) mutant, wild type and common peptides. Intensity is measured as counts per second (CPS) and the amount of each peptide quantified by this method is shown in fmols (inset box).

**Figure S2 related to Figure 1D**. Normalized V5 tag counts over time using unmapped reads.

**Figure S3 related to Figure 1**. Box plot of *FOXL2* expression across AGCTs using the RNA-seq dataset from Shah et al.^7^ after re-processing with GRCh37 and Ensembl annotations v69 (Table S1).

**Figure S4 related to Figure 2.** Pie charts showing genomic annotation of all FOXL2^WT^ (A) and FOXL2^C134W^ (B) derived ChIP-seq ERs using ChIPseeker R package^59^. Only peaks located on autosomes and sex chromosomes and not overlapping any ER from Dox empty vector samples were included. Promoters were defined as transcription start site +/- 2kb.

**Figure S5 related to Figure 2E**. Heat maps of RPKM values associated in Dox MUT (4 replicates) and Dox WT (3 replicates) and Dox EV (3 replicates) with statistically called FOXL2 ERs. Two kilobases on each side of the ERs are shown. Heat maps are ordered based on the specificity of the FOXL2 ERs: mutant-specific ERs and ERs shared between wildtype and mutant (minimum 50% overlap reciprocally between Dox WT and Dox MUT). Dox WT and Dox MUT ERs were defined as not overlapping with ERs found in Dox EV.

**Figure S6 related to Figure 2F**. Histogram of motif density in the mutant-specific enriched regions (A) and in the shared peaks (B) using the motif from the *de novo* motif analysis and represented below the x-axis.

**Figure S7 related to Figure 4.** EMSA with positive control CYP19A1 (aromatase PII promoter) probes from Fleming at al ^12^. Probe sequence is

5’ TTTCTTGGGCTTCCT**TGTTT**TGACTTGTAACCATAAAT 3’. Mutated probe sequence is `5’` TTTCTTGGGCTTCCT**TG**aggTGACTTGTAACCATAAAT 3’.

## Supplementary Tables Legends

**Table S1**. RPKM matrix for protein coding genes generated from AGCT and SVOG3e RNA-seq (polyA and ribodepleted).

**Table S2.** Genes differentially expressed in induced wildtype and mutant compared to induced empty vector control after 12h Dox induction (FDR 0.05). Only genes up- or down-regulated in a minimum 3 out of the 4 induced mutant replicates, and in a minimum 2 out of the 3 induced wildtype replicates were included. Genes already known to be FOXL2 targets are indicated with the reference paper.

**Table S3**. Enriched known motifs in the shared peaks (147,094) (first spreadsheet: ‘SHARED’) and in the mutant-specific peaks (61,564) (second spreadsheet: ‘C134W_specific’).

