## Supplemental figures 1 to 7 for "The pathognomonic FOXL2 C134W mutation alters DNA binding specificity"

### Heavy Peptide FOXL2 standard Curve

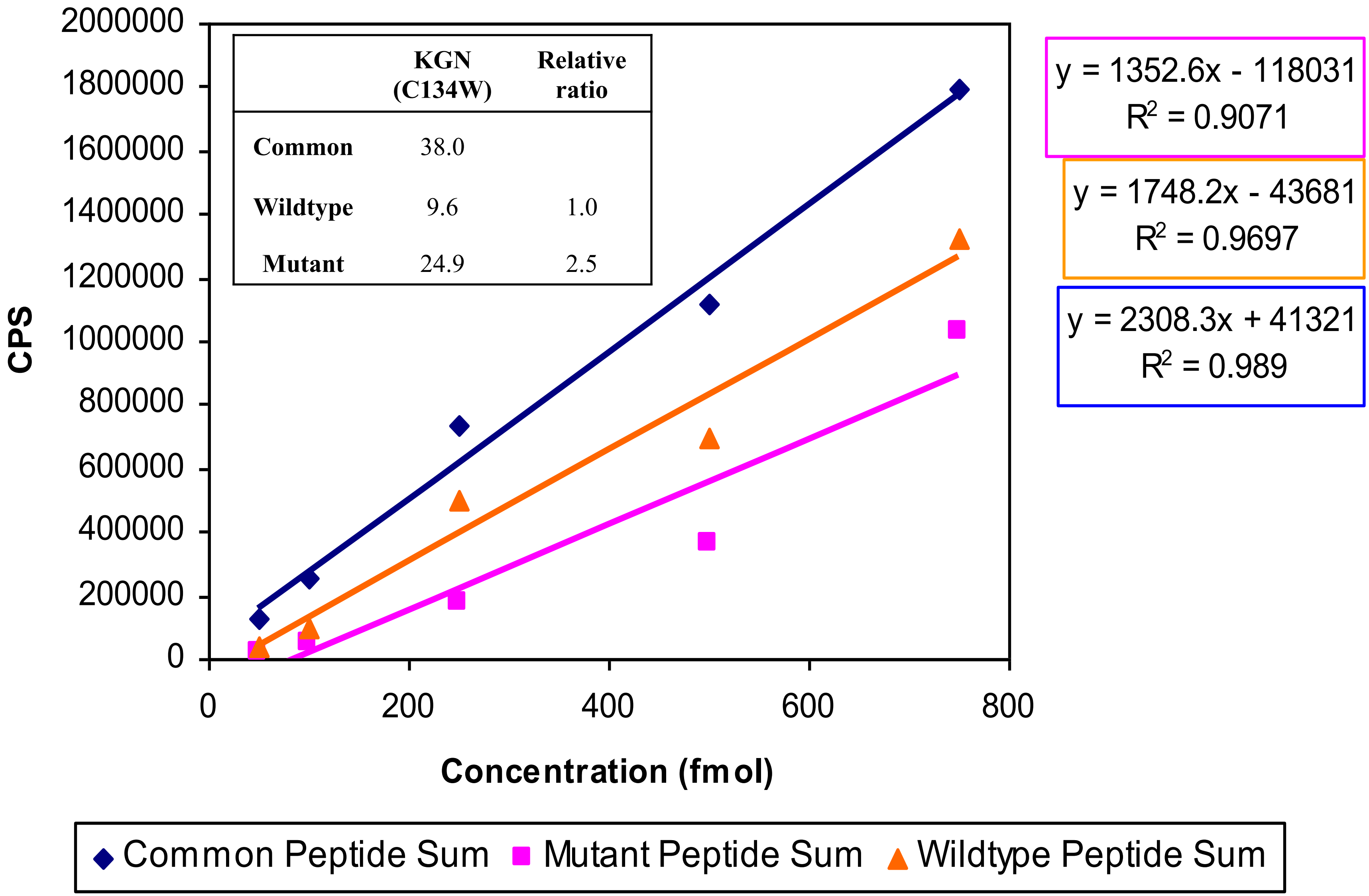


| | KGN (C134W) | Relative ratio |
|---|---|---|
| Common | 38.0 | |
| Wildtype | 9.6 | 1.0 |
| Mutant | 24.9 | 2.5 |

Fig. S2

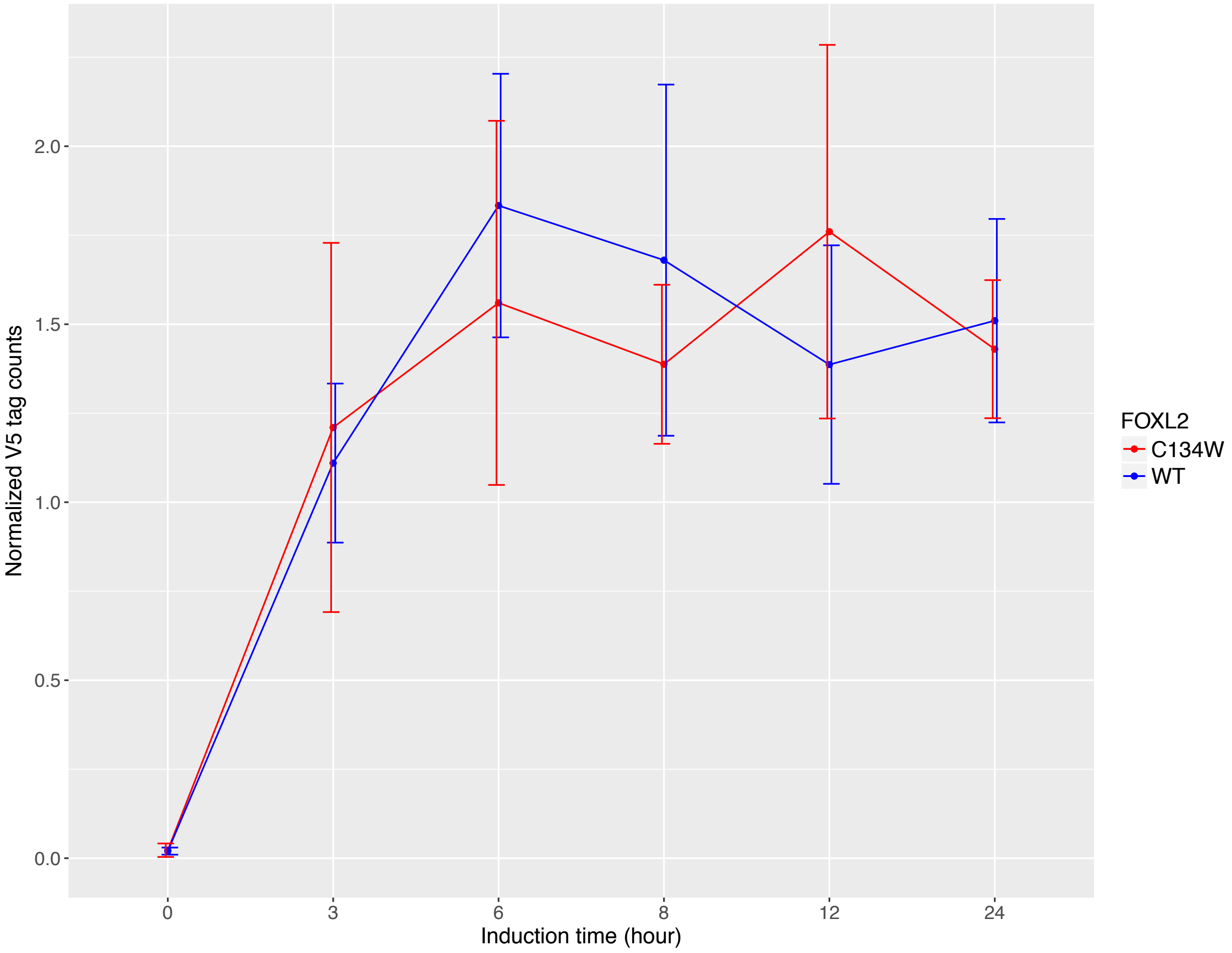

Normalized V5 tag counts
0.0
0.5
1.0
1.5
2.0
0
3
6
8
12
24
Induction time (hour)
FOXL2
C134W
WT

Fig. S3

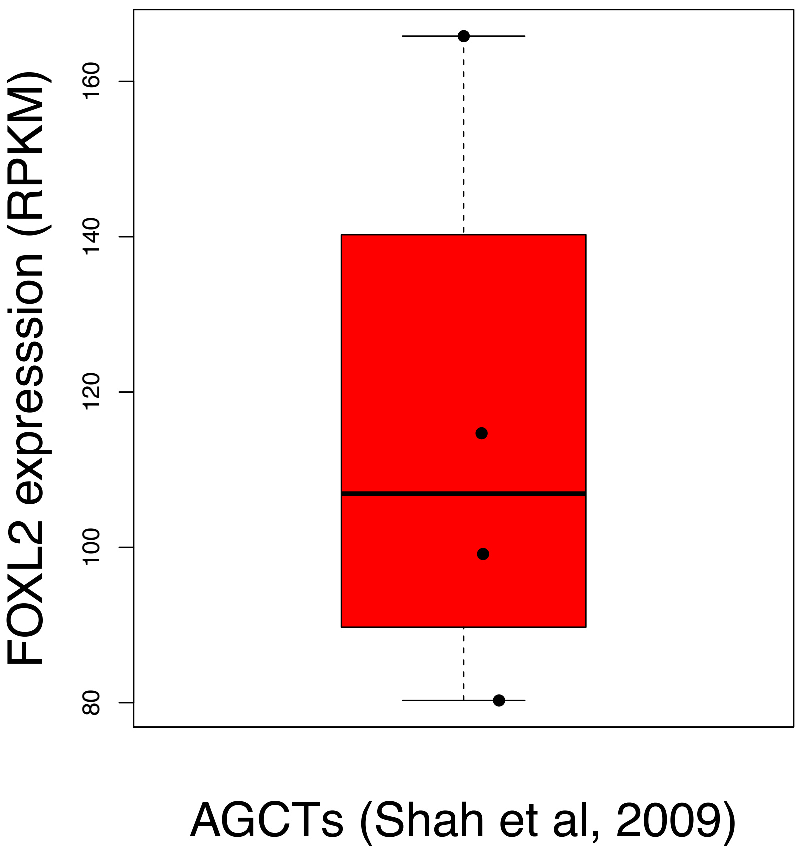
FOXL2 expresssion (RPKM)
80
100
120
140
160
AGCTs (Shah et al, 2009)

Fig. S4

A

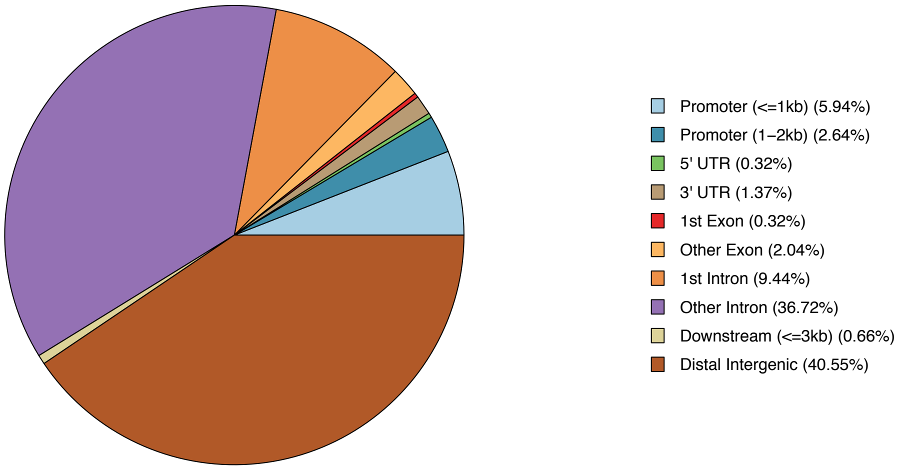

Promoter (<=1kb) (5.94%)
Promoter (1–2kb) (2.64%)
5' UTR (0.32%)
3' UTR (1.37%)
1st Exon (0.32%)
Other Exon (2.04%)
1st Intron (9.44%)
Other Intron (36.72%)
Downstream (<=3kb) (0.66%)
Distal Intergenic (40.55%)


B

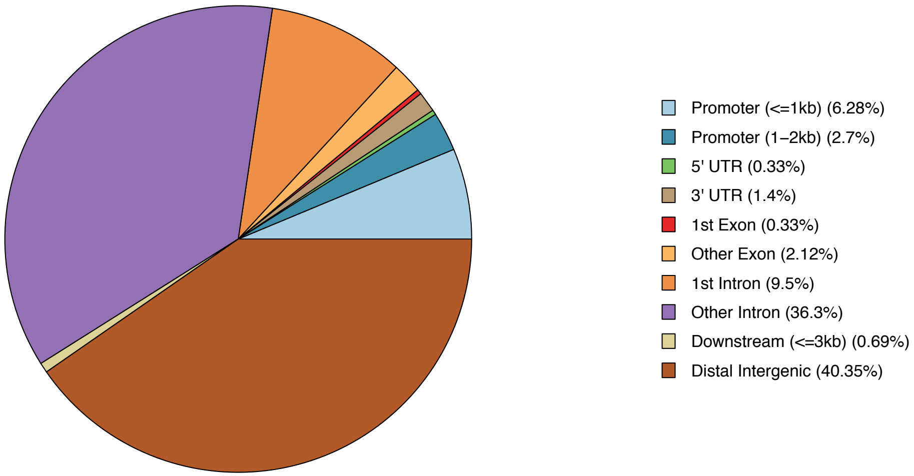

Promoter (<=1kb) (6.28%)
Promoter (1–2kb) (2.7%)
5' UTR (0.33%)
3' UTR (1.4%)
1st Exon (0.33%)
Other Exon (2.12%)
1st Intron (9.5%)
Other Intron (36.3%)
Downstream (<=3kb) (0.69%)
Distal Intergenic (40.35%)

Fig. S5

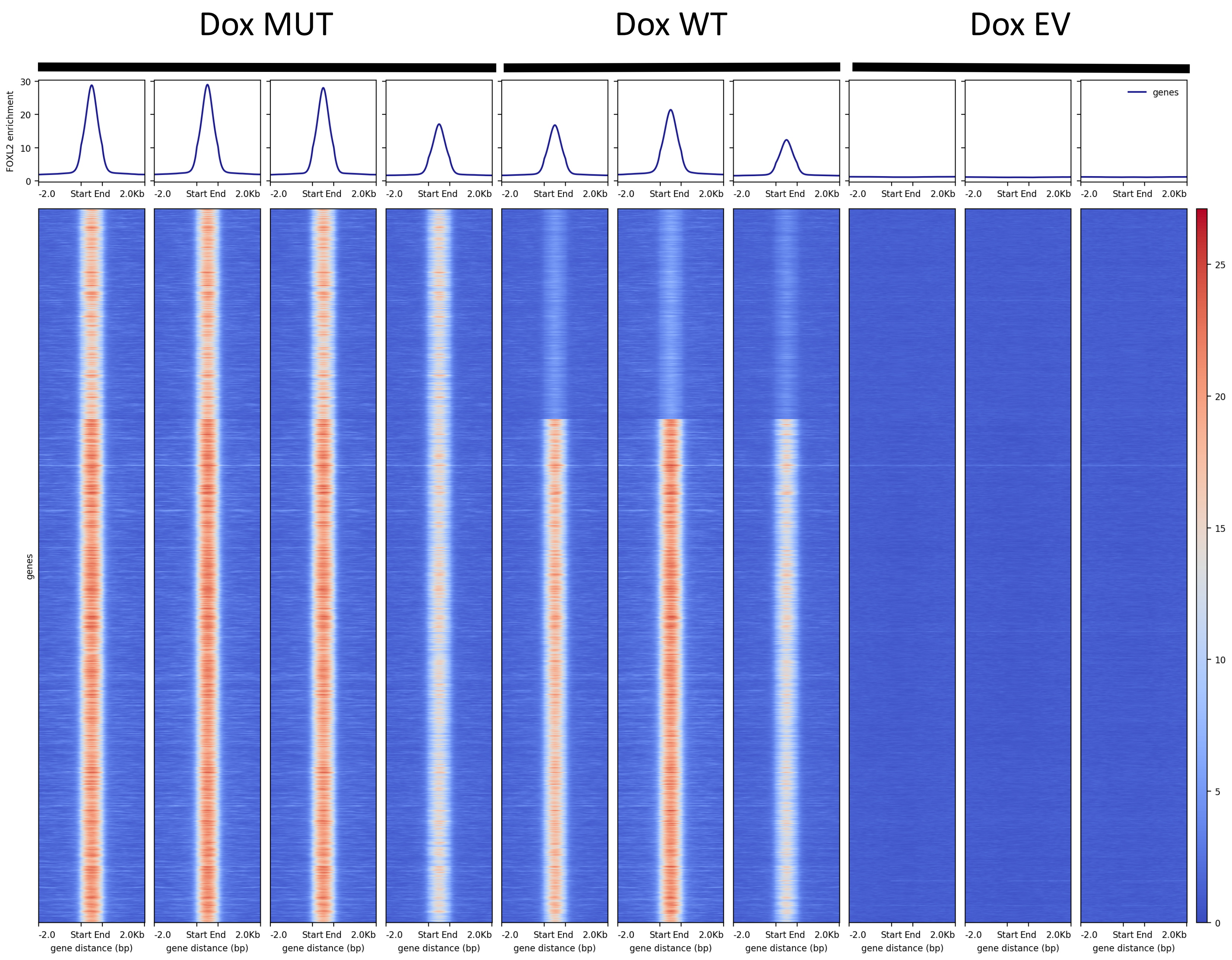

Dox MUT
Dox WT
Dox EV
FOXL2 enrichment
genes
-2.0
Start End
2.0Kb
gene distance (bp)

Fig. S6

A

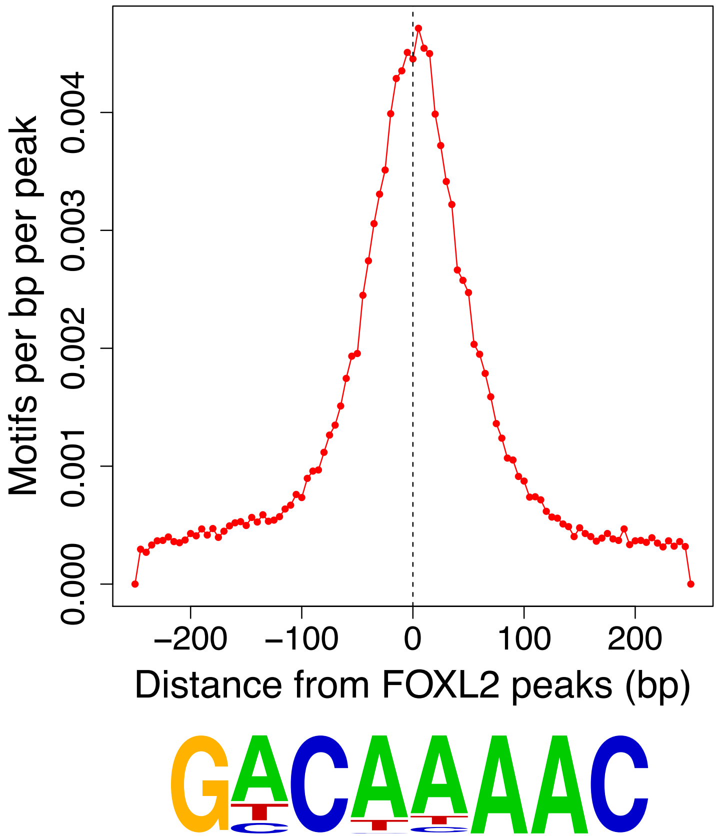

Motifs per bp per peak
0.000
0.001
0.002
0.003
0.004
−200
−100
0
100
200
Distance from FOXL2 peaks (bp)
GACAAAAC


B

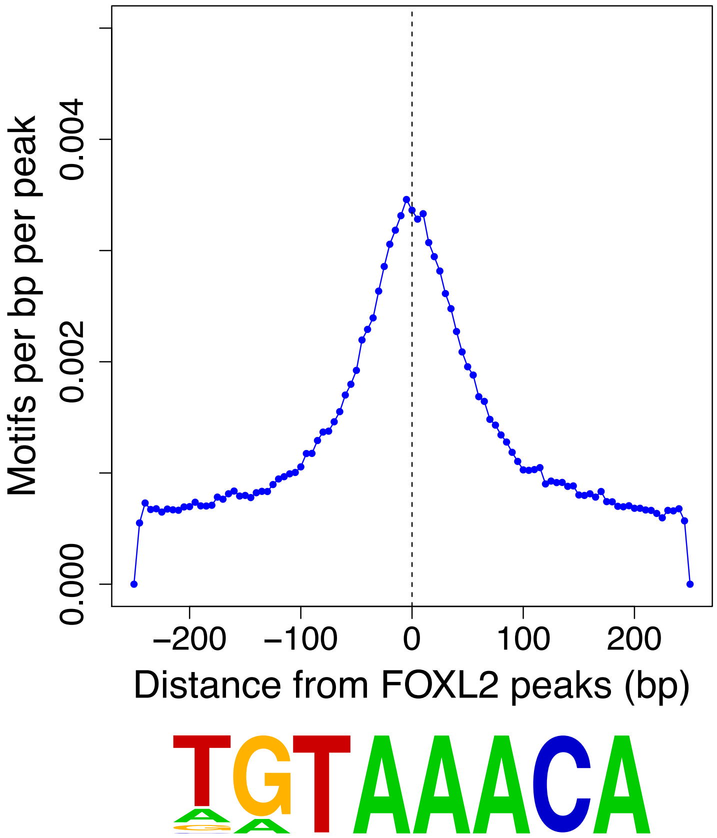

Motifs per bp per peak
0.000
0.002
0.004
−200
−100
0
100
200
Distance from FOXL2 peaks (bp)
TGTAAACA

Fig. S7

| | | | | | | | |
|---|---|---|---|---|---|---|---|
| IR labeled probe | + | + | + | + | + | + | + |
| FOXL2 WT | | + | | + | | + | |
| FOXL2 C134W | | | + | | + | | + |
| Competitor (1:2) | | | | + | + | | |
| Mutated competitor | | | | | | + | + |
| Lane | 1 | 2 | 3 | 4 | 5 | 6 | 7 |

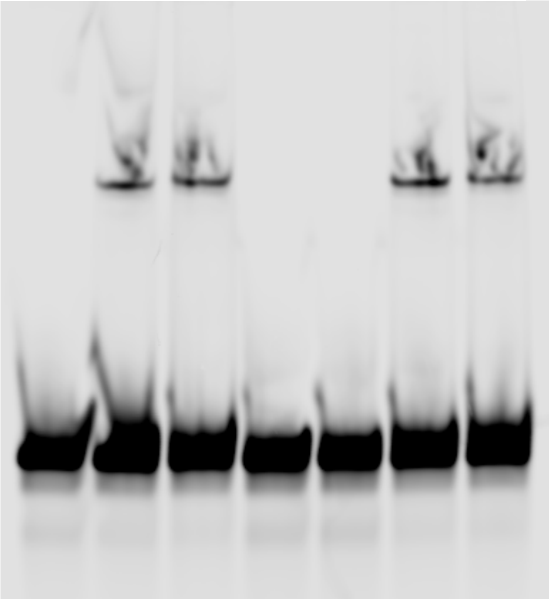
